# UV reflectance in crop remote sensing: Assessing the current state of knowledge and extending research with strawberry cultivars

**DOI:** 10.1101/2023.05.04.539478

**Authors:** Megan Heath, David St-Onge, Robert Hausler

## Abstract

Remote sensing of spectral reflectance is a crucial parameter in precision agriculture. In particular, the visual color produced from reflected light can be used to determine plant health (VIS-IR) or attract pollinators (Near-UV). However, the UV spectral reflectance studies largely focus on non-crop plants, even though they provide essential information for plant-pollinator interactions.

This literature review presents an overview of UV-reflectance in crops, identifies gaps in the literature, and contributes new data based on strawberry cultivars. The study found that most crop spectral reflectance studies relied on lab-based methodologies and examined a wide spectral range (Near UV to IR). Moreover, the plant family distribution largely mirrored global food market trends.

Through a spectral comparison of white flowering strawberry cultivars, this study discovered visual differences for pollinators in the Near UV and Blue ranges. The variation in pollinator visibility within strawberry cultivars underscores the importance of considering UV spectral reflectance when developing new crop breeding lines and managing pollinator preferences in agricultural fields.

## Introduction

Precision agriculture is a modern farming approach that aims to optimize crop production by using advanced technologies and data analysis techniques. By incorporating modern technology with traditional farming principles, farmers can now manage fields with minimal inputs and human resources. One of the key tools used in precision agriculture is remote sensing, which is based on electromagnetic radiation (Wójtowicz M. et al.,2016; Xue, J., & Su, B., 2017). Remote sensing is used to capture data from vegetal surfaces to generate field maps that can be used to characterize biophysical features such as water and nutrient stress, presence of infection/disease or overall growth of crops.

To capture data from vegetal surfaces, remote sensors are used to capture electromagnetic reflectance in three key spectral regions (Xue, J., & Su, B. 2017) including ultraviolet (UV), visible and near-infrared (IR). This data is then analyzed using vegetation indices (VI) to derive insights into the health, growth and overall well-being of crops (Zhang & Kovacs, 2012; Liu C. et al., 2016).

With these insights, farmers can adjust their crop management strategies in real-time to optimize crop yields, reduce waste and increase profits. This has led to the development of new techniques for crop management, such as precision irrigation, soil mapping and crop yield forecasting, all of which are crucial for improving efficiency and reducing waste in modern agriculture.

### Flower Patterns in UV

Since Darwin’s time (1876), researchers studying pollinator interactions have focused on the diverse colors and patterns of flowers. Pollinating species, unlike many foraging predators, have receptors for UV (Briscoe & Chittka, 2001), leading plants to develop UV floral patterns to attract beneficial insects visually while remaining cryptic to foraging species. Compounds that absorb or reflect radiation are arranged in patterns on reproductive structures like anthers and petals, signaling feeding locations and differentiating plants from con-specifics even at the cultivar level (Yoshioka et al., 2005).

Size, shape, and contrast can influence the visibility of these patterns, particularly from the air, and affect how visible a plant is to its con-specifics. Floral signaling strategies may respond to the perceptual constraints of pollinators, as Spaethe et al. (2001) observed that bumblebees favored high color contrast on the floral surface for large flowers (e.g., UV pattern) but only favored high contrast with green foliage for small flowers. This resulted in optimal foraging strategies and more accurate floral recognition while in flight.

A typical signaling pattern of large flowers is the ”bullseye” pattern, where flowers consist of UV-absorbing centers and UV-reflecting peripheries. Previous studies found that bees make their first antennae contact with the UV-absorbing part and untrained bees preferentially visit bullseye-patterned flowers (e.g., Koski Ashman, 2014; Papiorek et al., 2016).

### Factors affecting crop visibility to pollinators

Factors affecting crop visibility in the UV spectrum are reducing UV light on crop surfaces and decreasing UV-reflective pigments due to breeding efforts. Uv reflective patterns become less visible under the physical conditions of greenhouses that commonly employ UV-blocking coverings (Morandin, L. A. et al. 2002). In a study by Morandin et al. (2001), four types of polyethylene greenhouse coverings, varying in their UV transmittance, found that bees made twice as many foraging trips under low UV transmittance plastics. Furthermore, 136 percent more bees remained within the greenhouse after ten days, drastically affecting operation costs and crop production.

Bee pollination of crops results in heavier, more uniform crops, which fetch a higher market value. Therefore, hives are often supplemented in agricultural settings (e.g. Klatt, B. K. et al. 2014). Another factor for commercial growers to consider is the genetic component of UV patterns when breeding new cultivars. Brock M. T. et al. (2016) showed that UV patterning varied greatly among Brassica rapa genotypes and that insects preferred flowers with UV patterns over those without patterns, such as their wild relatives. Moyers et al. (2017) found that the UV pattern of sunflowers could be modified without affecting flower head size based on the mapped genetic architecture. Flower head size is a critical trait for breeding this crop and could have unintended effects on pollinator–flower interactions. Breeders need to consider the genetic architecture of a crop when creating new cultivars. Research supports that colour patterning in various crop families varies significantly with heredity (e.g., Mangelsdorf, A. J., & East, E. M.,1927; Henz, A. et al., 2015; Muchhala, N. et al., 2014). However, few studies have explored the use of UV floral reflectance of plants, even fewer for crops specifically. Spectral reflectance studies have reported down to 300 nm, but most species in reflectance databases (e.g., FReD) are native species, not crop species (Arnold S. et al. 2008).

### UV floral reflectance of Rosacea crops

The Rosacea family includes several common orchard and berry crops like apples, cherries, raspberries, and strawberries, making it one of the four primary crop families grown commercially in greenhouses worldwide (Guerra-Sanz, 2008). Notably, members of the Rosacea family, including blackberry and almond cultivars, display consistent and distinct peaks in the near UV range, suggesting a potential role of UV patterning in pollinator signaling within the family (Gyan & Woodell, 1987; Chen et al., 2019).

Strawberries are an extensively cultivated crop used widely in both greenhouses and traditional fields worldwide. The global greenhouse production alone was a staggering $34.8 billion industry in 2021, with North America holding the largest market share of 32.8 percent (Precedence Research, 2022). In terms of total global fruit production, strawberries and tomatoes were the two leading crops, representing 25.4 percent and 56.7 percent of total vegetable production, respectively (Forbes, 2022). Notably, *Rosacea* crops, with strawberries comprising 75.4 percent of berry crops, constituted the majority of global fruit production in 2021 (Statista, 2023), competing with the likes of *Musaceae*(bananas and plantains), *Rutaceae* (citrus), and *Cucurbitaceae* (melon) families.

Despite the widespread cultivation of strawberries, research on the UV floral reflectance of strawberry cultivars remains scarce. Though most strawberry flowers appear white to human eyes, Ceuppens et al. (2015) found differences in pollination of two related strawberry varieties when cultivated together, potentially due to discrepancies in floral patterning rather than the presence of volatile floral substances. Thus, UV floral reflectance differences may be a relevant factor here. Notably, there is a lack of systematic literature reviews on UV floral reflection of crop species, which this present study aims to address by documenting the current state of floral UV-reflectance of crops in scientific literature and expanding on it with strawberry cultivars.

## UV crop reflectance meta-analysis

### Literature review Methodology

We followed the eight-step guide to conducting a meta-analysis by Hansen, C. et al. (2021) in this literature review. We analyzed scientific articles studying the UV-reflectance of crops. We searched the electronic database Scopus (1969-2020) for the following keywords: UV* OR ultraviolet AND camera AND/OR ”Spectral reflectance” AND ”flo*” AND/OR ”crop*” AND/OR ”plant*”. In addition, searches were limited to the English language, publication in a journal or conference proceeding, and fell within the categories ¡agriculture¿, ¡botany¿, AND/OR ¡environmental sciences¿. A total of 1013 articles met the search criteria and were screened for crop species and spectral reflectance under 400 nm using a single reviewer. We excluded from this review papers that dealt with the spectral reflectance of compounds derived from plants in chemical isolation or studied UV spectral fluorescence rather than reflectance. UV Reflectance measures a reflected wavelength in the near UV range (300-400 nm), often used by flowering plants for pollinator signalling. In contrast, UV fluorescence is a visible emission of wavelengths due to a substance or pigment’s absorbance of UV radiation. Until recently, the terms were used interchangeably in literature; therefore, we carefully examined the methodologies employed. In total, 170 papers related to botanical plants, of which 149 covered spectral reflectances below 400 nm in some capacity. When filtered for agriculturally relevant species, 52 records remained from 29 families and 73 crop species, as listed in Tab. 1.

**Table 1.**
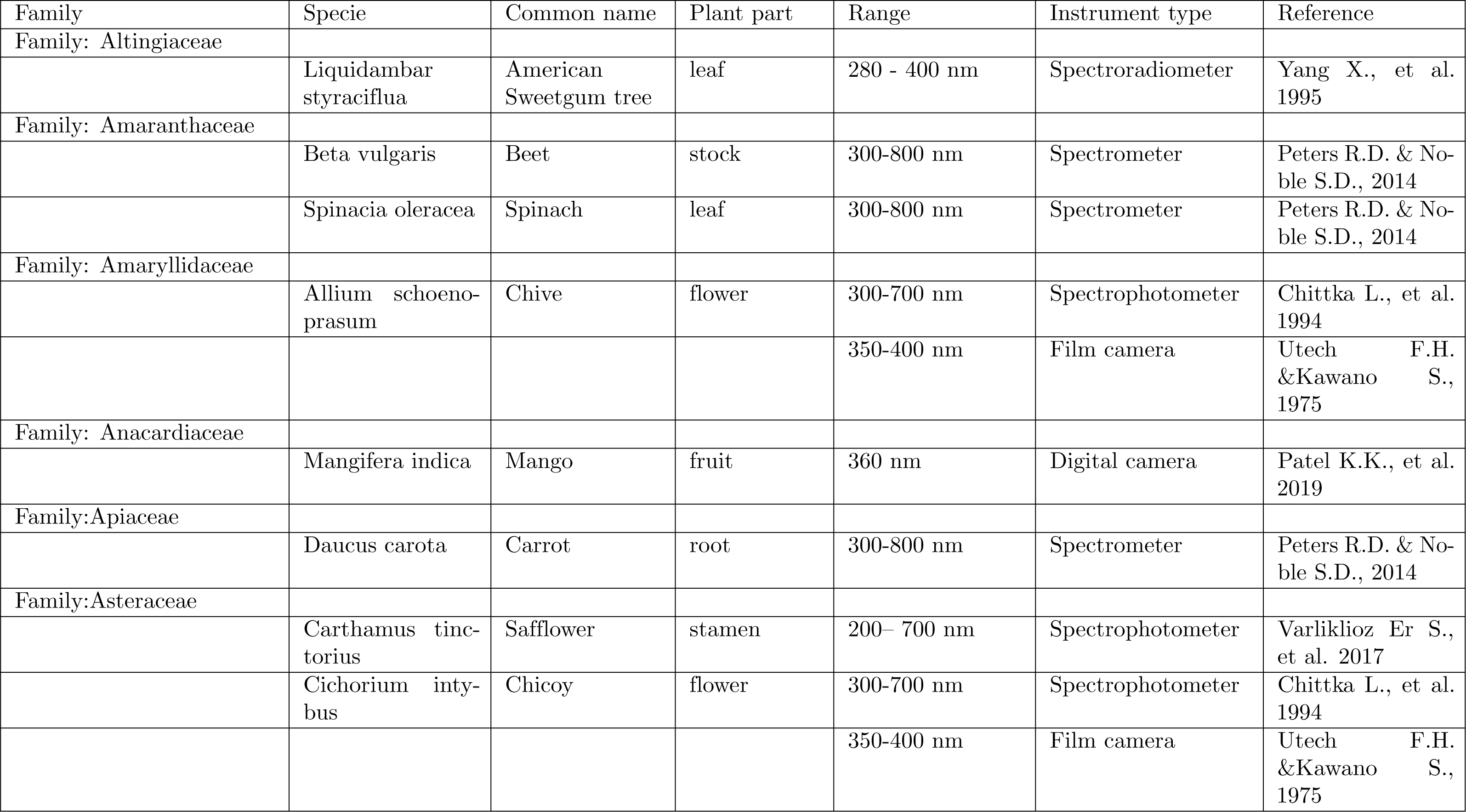
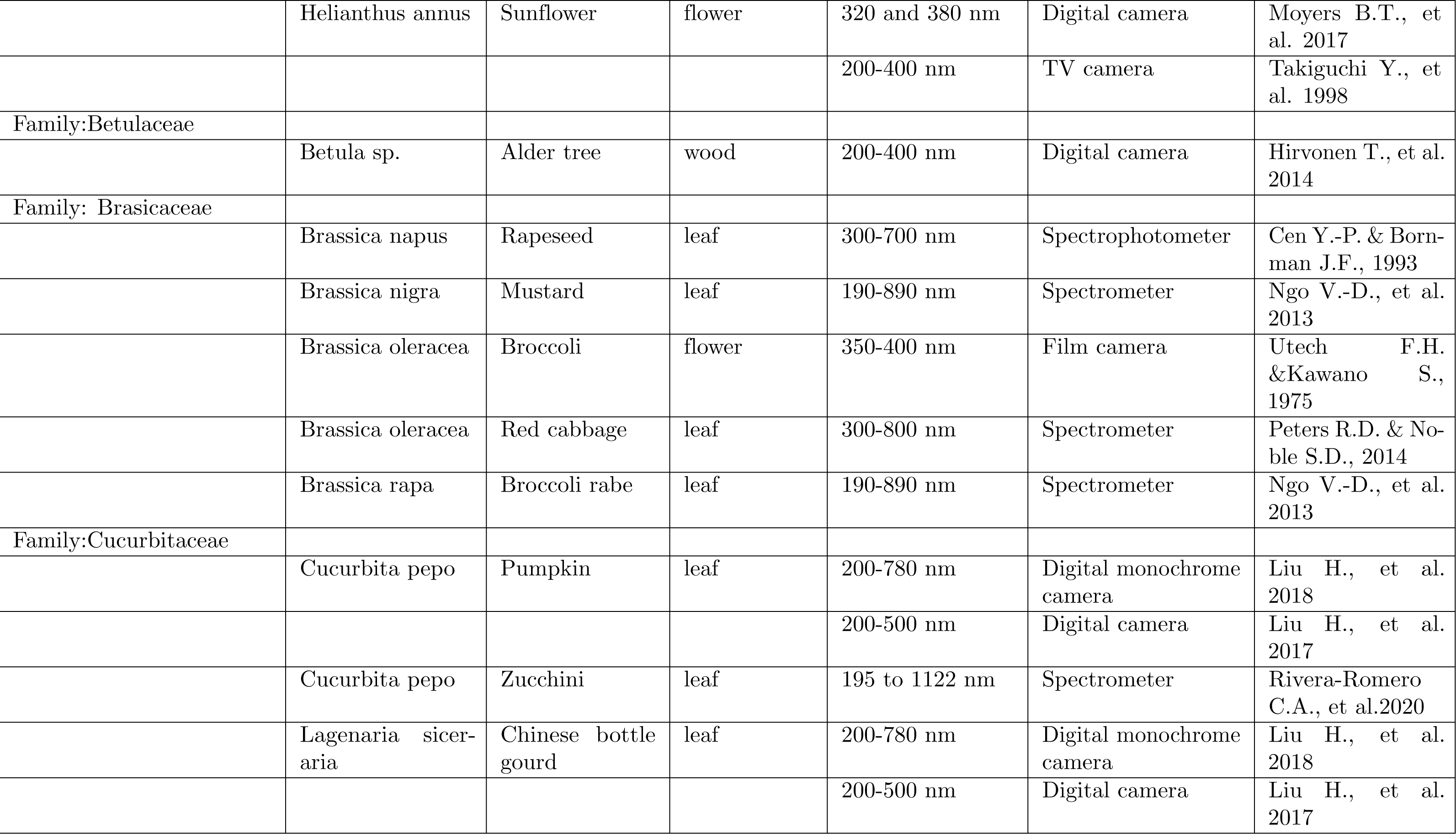
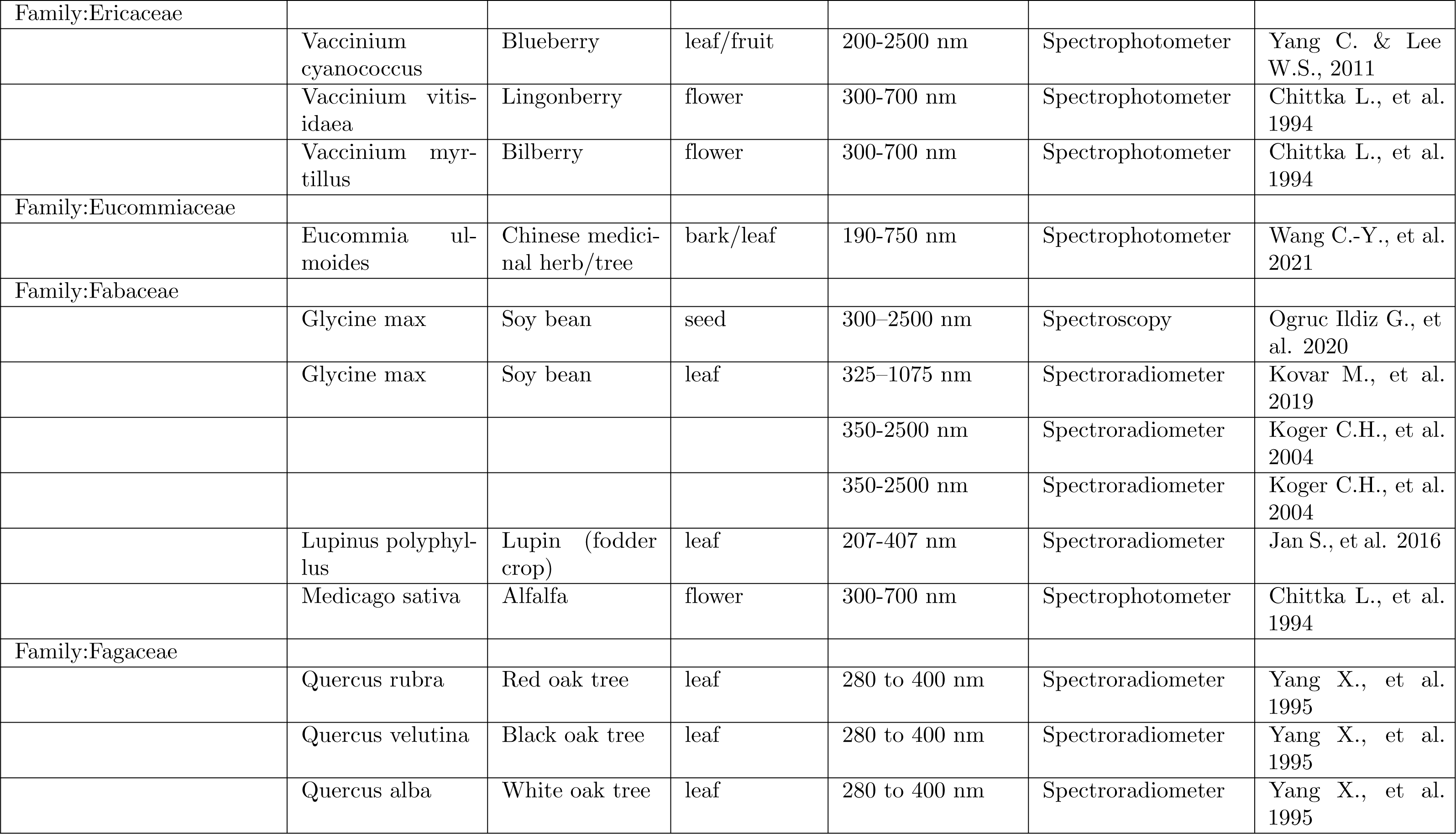
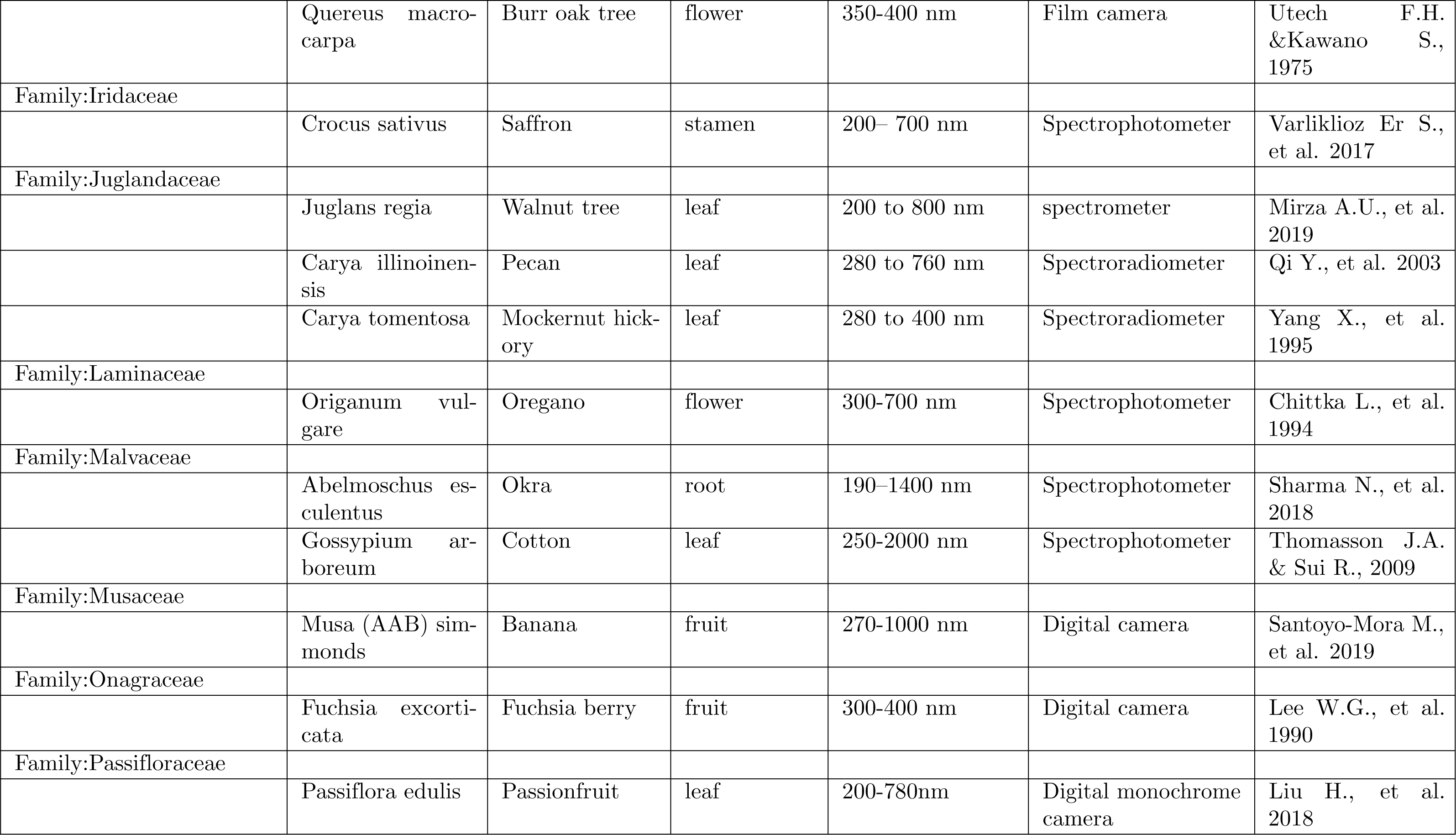
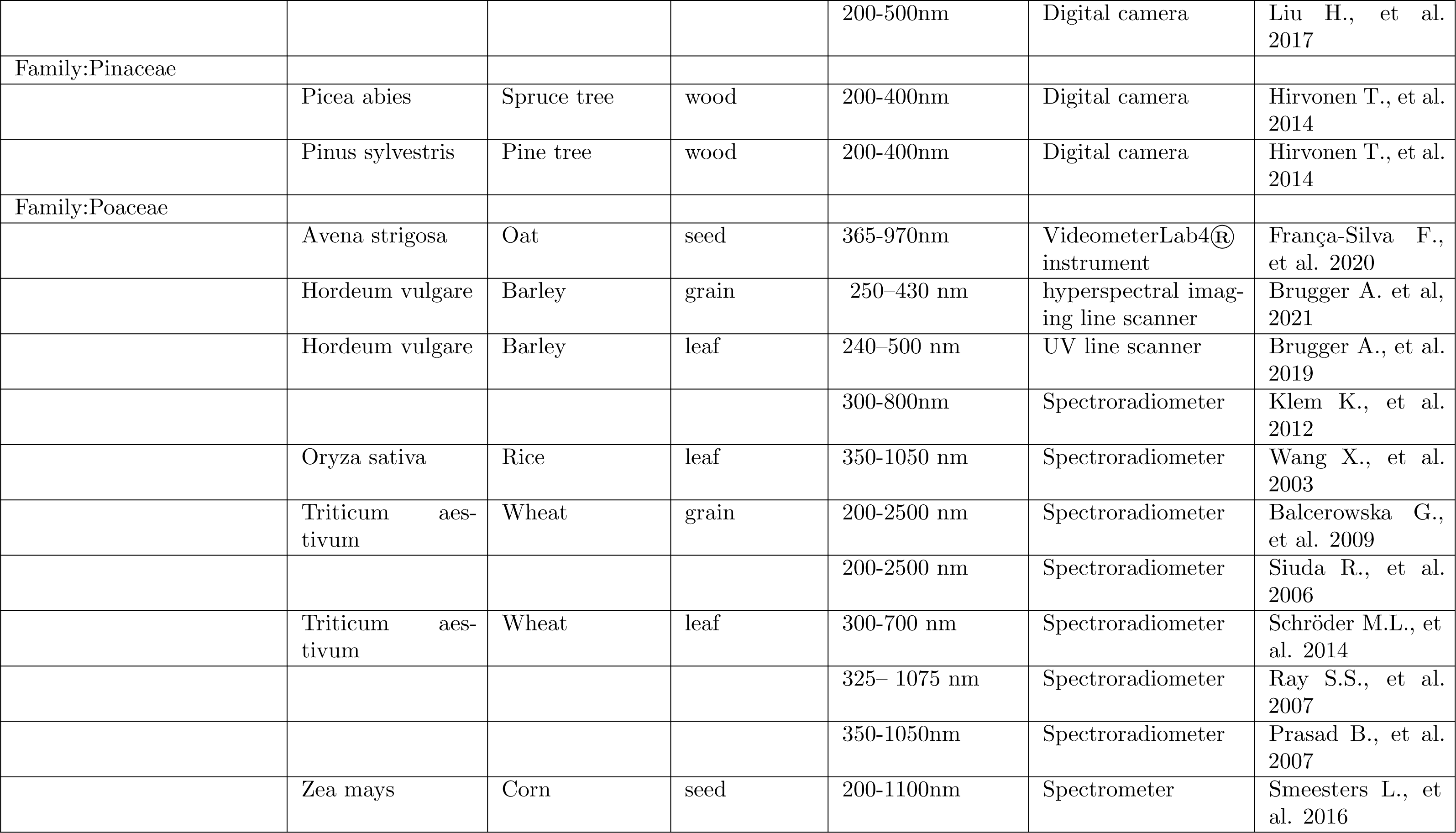
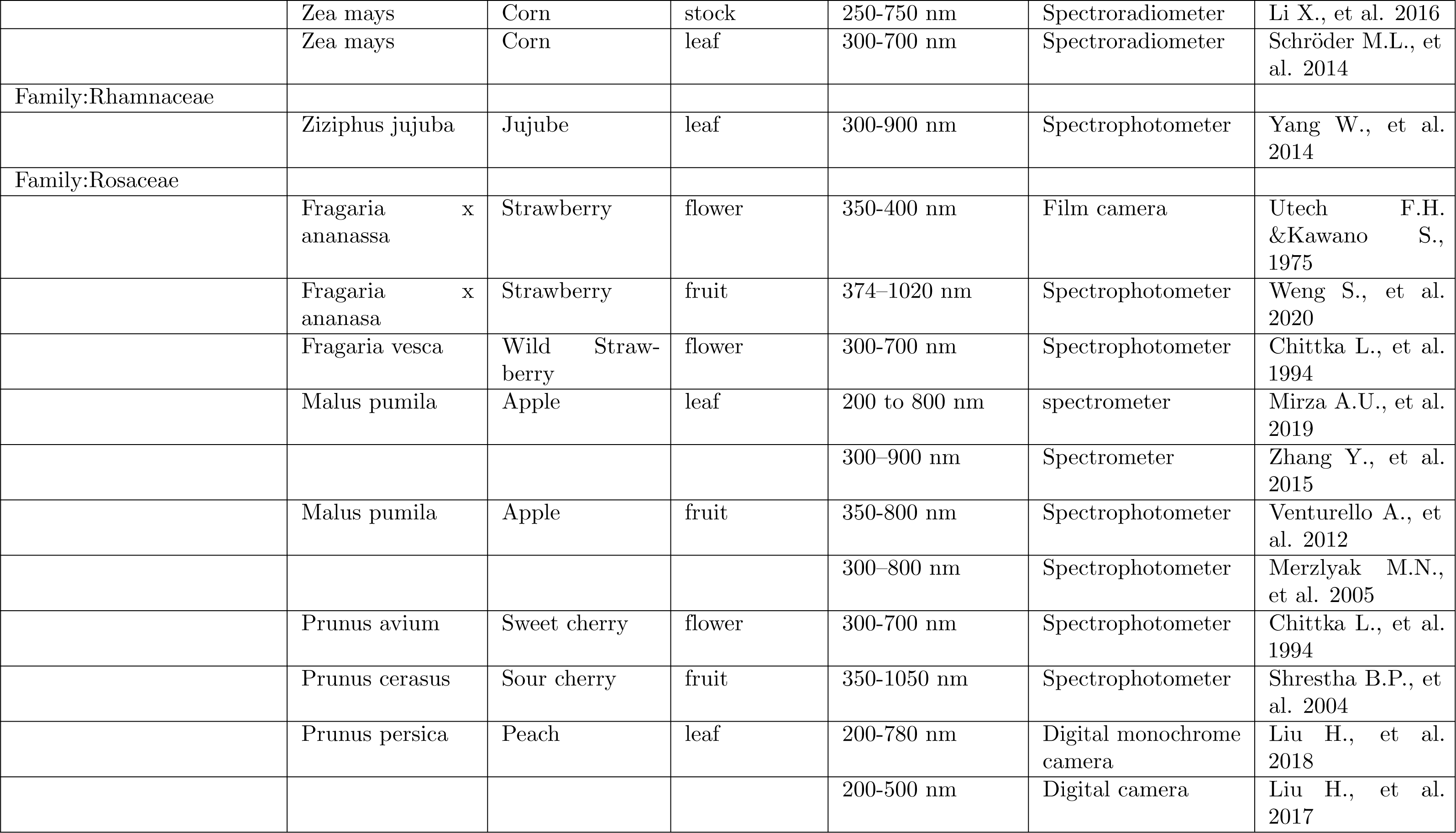
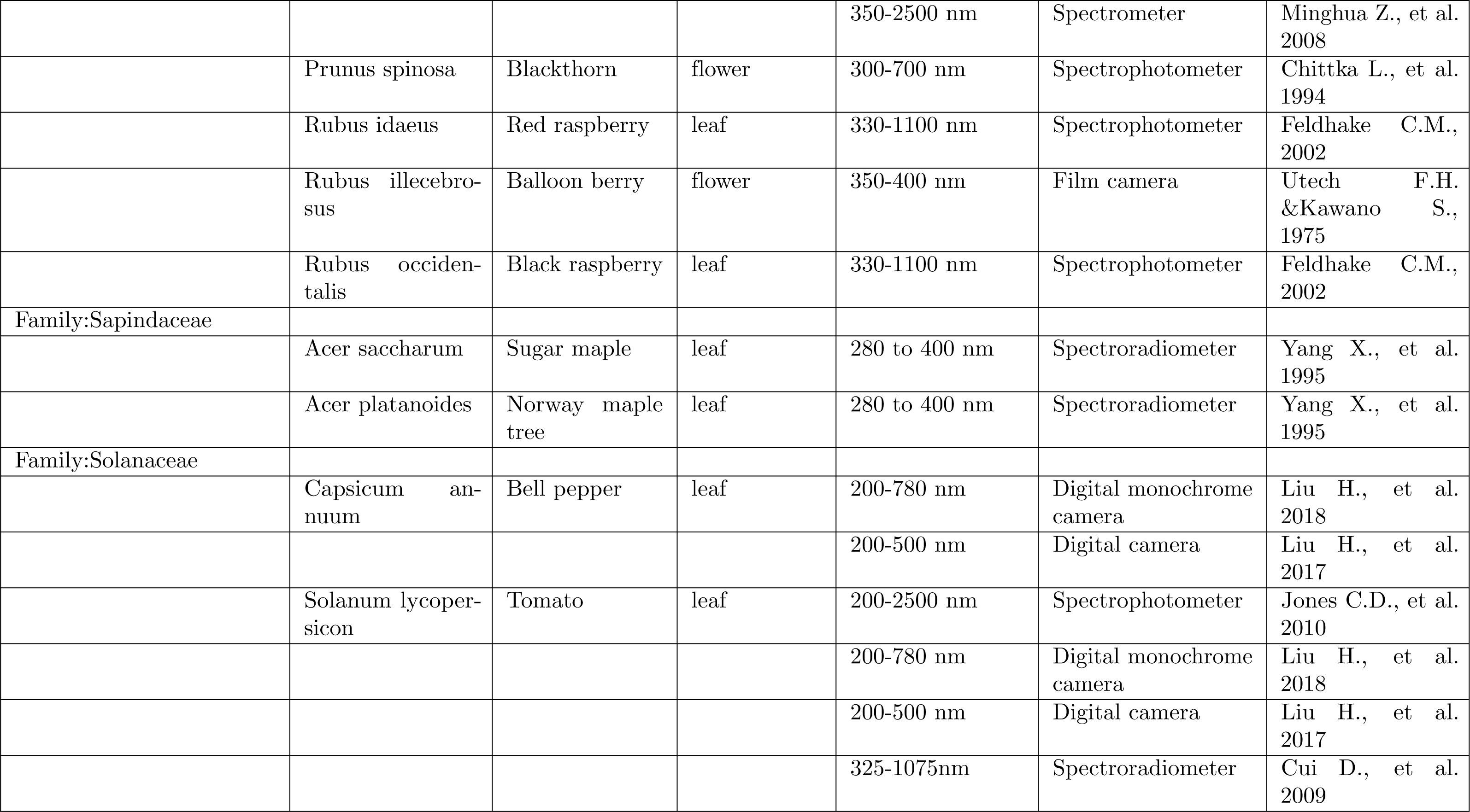
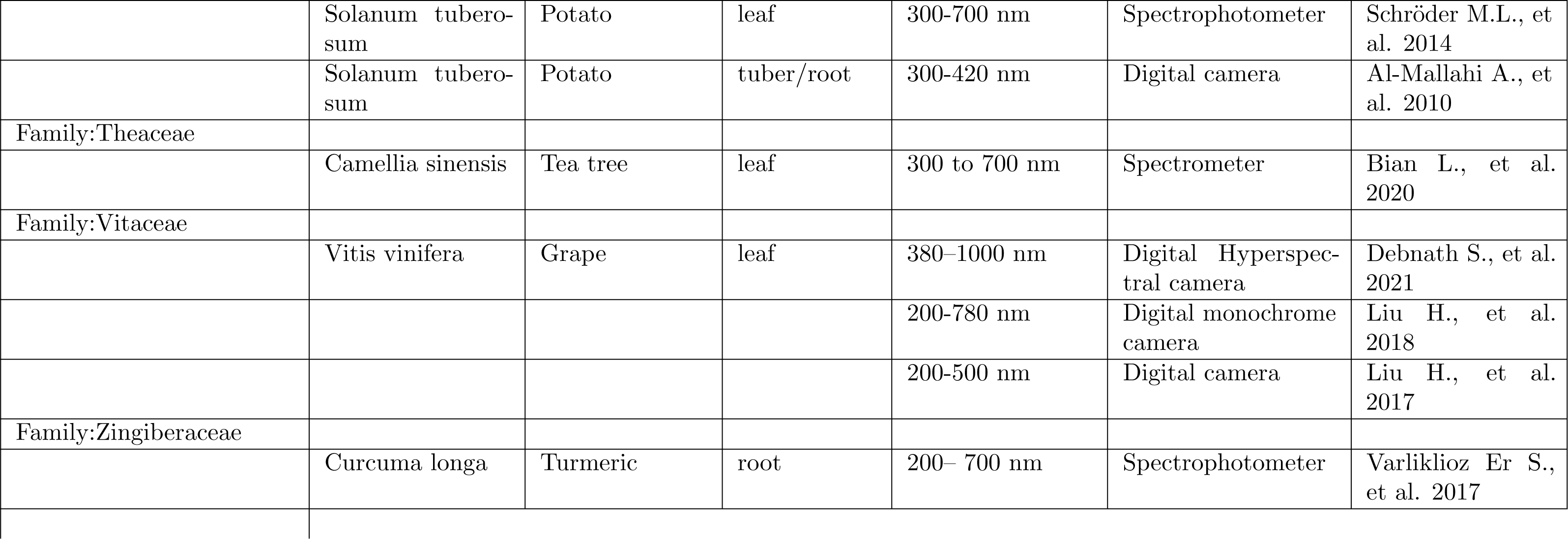
Major characteristics of studies included in the meta-analysis from 1969-2020

#### Metrics

Articles meeting the above criteria had the following parameters noted: instrument model used, the spectral range for measurements, floral part and species analyzed, and year of publication.

Instrument models were grouped into 4 categories: Camera, Videometer, spectrometer/spectroradiometer, and spectrophotometer. Cameras were defined as self-contained, image-recording devices which relied on an external light source. This included video, monochrome, multi-spectral, and hyper-spectral cameras which employed CMOS or CCD sensors, as well as UV film cameras. Spectrometer and spectroradiometer were grouped together as the terms are often used interchangeably. A spectrometer measures the reflectance spectrum of an object or substance. Its sensor array can separate out the light received at each wavelength and generate an amplitude graph of the incoming signal. A spectroradiometer can also take calibrated readings of power, intensity, and radiance of the incoming signal at each wavelength (International Light Technologies Inc.,2019). On the other hand, Spectrophotometers measure the light absorption or transmission of a sample. A reflectance curve can then be generated from the absorption and transmission measurements using Kirchoff’s law (Spectrecology,2021).

Videometer was its own category as it utilizes an integrating sphere with a light-emitting diode, similar to a spectrometer; however, the sensor captures a pixelated image of an object at each wavelength (Carstensen, J. M.,2022). For all instruments, spectral ranges were binned according to the following nanometer (nm) ranges: near UV (300-380nm), Blue (381-520nm), Green (521-625nm), red/ IR (*>*625nm) in accordance with the international society for optics and photonics (Malacara, D., 2011).

Floral parts analyzed were grouped into 5 categories: flower, stem, leaf, fruit, and root. Flower included the anther, stamen, petal, and sepal elements of a plant’s reproductive structure. The stem encompassed dermal (cork & bark), vascular (xylem & phloem), and ground tissues (parenchyma, collenchyma,& sclerenchyma). Fruit encompassed seed and/or ripened ovary of a flowering plant. Root included tubers as well as roots themselves. Leaf category contained upper and lower sides of leaves. The species studied were divided by plant family to assess trends in the literature.

Trends in research over time were assessed by cross-referencing the above parameters with the publication year.

## Results

### Instrumentation

Most methodologies consisted of lab bench setups due to the size, weight, and equipment cost (e.g., Spectrophotometer). Of the data collection methods in Tab. 1, less than a third (31.7%) used cameras. UV Film represented 20% of data collected and occurred before 1980. After 1980 digital data collection using cameras became standard. The average cost of the cameras was $772.50 CAN and varied in weight from 50g to 2.75 kg, averaging 302g across all species recorded. Of note: no studies performed aerial remote sensing of UV reflectance. Fig. 1 depicts the trends in instrumentation.

**Fig 1.**
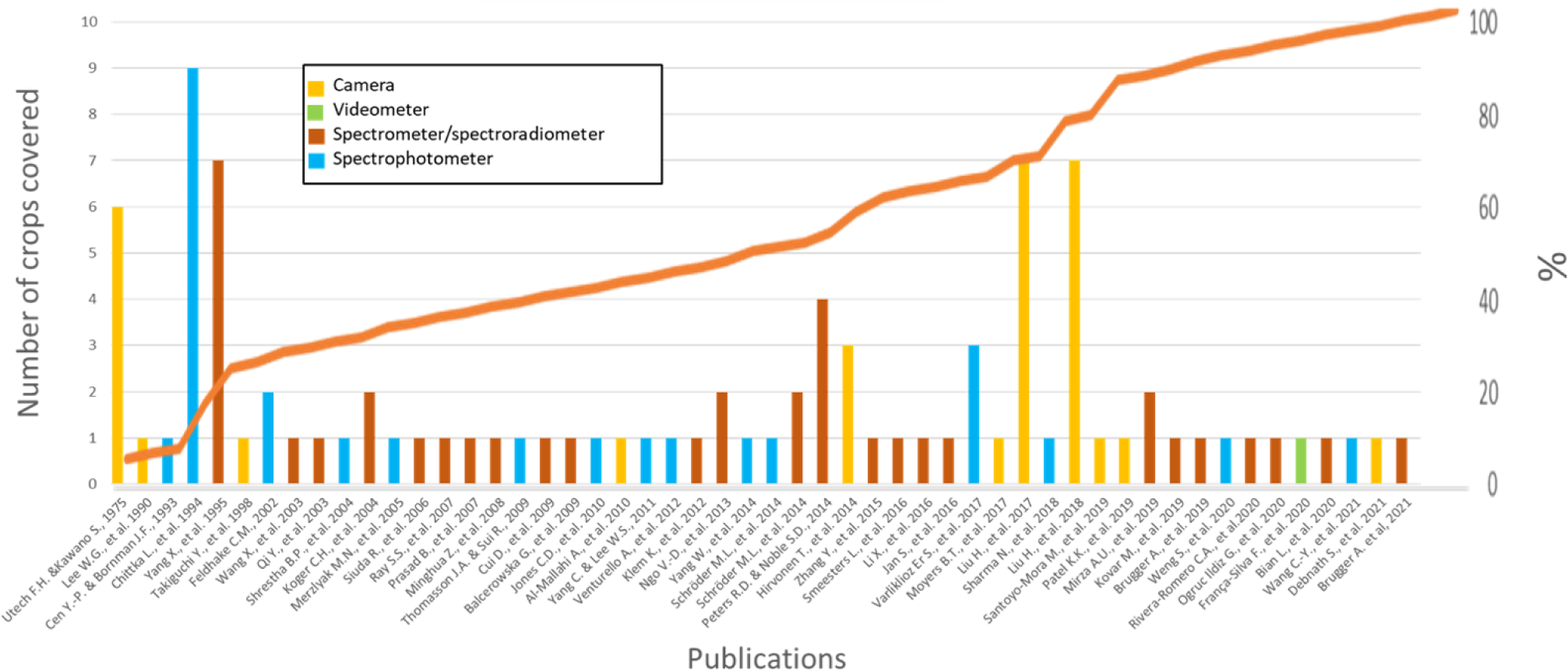
Trends in UV reflectance. Publications from meta-analysis ranging from 1960 to 2021 for crop UV spectral reflectance.

Spectrometers/Spectroradiometers and Spectrophotometers comprise the bulk of collection methods (36.8% and 27.4%, respectively) and have occurred consistently since the early 1990s. These are lab-based, costly devices, often shared with other departments, such as chemistry. They allowed for quantitative analysis using spectrograms compared to qualitative analysis with film cameras, as seen in Utech F.H. & Kawano S. (1975).

#### Spectral range

Though all publications in this study had to include the near UV range (300-380nm), many also presented visible and near IR spectrums. Cameras presented narrower ranges (Near Uv to blue) more often than any other instrument category, followed by Spectrometer /Spectrophotometers (Fig. 2). We attribute this disparity to the nature of the instrumentation chosen for the study. Digital cameras have a sensor that is more sensitive to Red and Near IR wavelengths. Therefore, a narrow range (usually Near UV to blue) must be captured using specialized lenses and filters to capture UV imagery. Comparatively, Spectrophotometers, Spectrometers, Spectroradiometers, and Videometers can capture data to the nanometer level without such interference.

**Fig 2.**
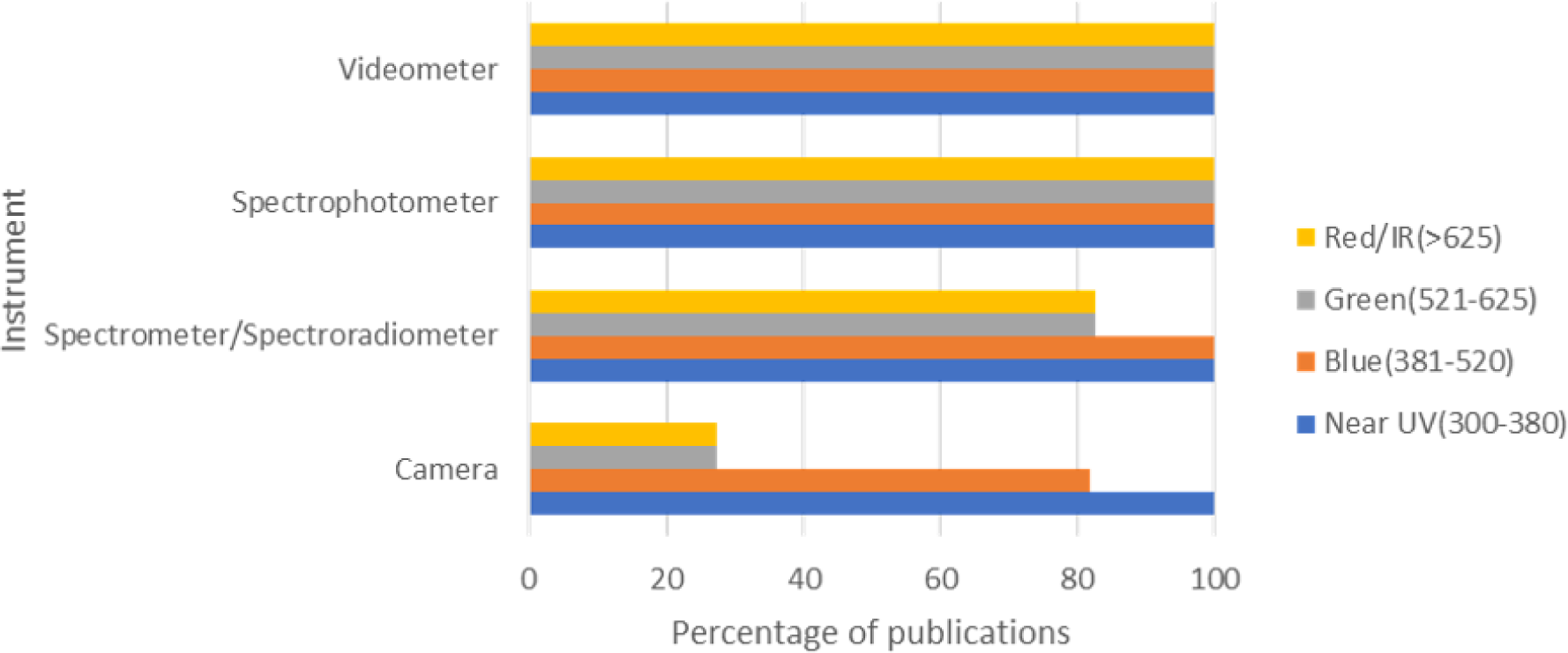
Spectral ranges presented in Crop UV reflectance literature.

Authors usually capture complete spectral ranges with these devices, even if the publication only interests a particular region.

#### Species and floral parts

As previously stated, 73 crop species from 29 families were included in this meta-analysis, and are listed in Tab. 1. Compared to the global production of fruits and vegetables in Fig. 3, we see an overlap in crop family representation from our meta-analysis in Fig. 4. Four of the top five families in our literature review (*Rosacea*, *Solanaceae*, *Fabaceae*, and *Brassicaceae*) overlapped with global production’s largest fruit and vegetable families in 2021. The *Rosaecea* family was the largest in global fruit production, whereas *Solanaceae*,*Fabaceae/Leguminaceae* and *Brassicaceae* were the top three vegetable-producing families globally in 2021, respectively. The above four families comprised 36.8% of the publications in our meta-analysis. The disproportional representation of the above families in our review supports that research decisions for crop species follow market trends.

**Fig 3.**
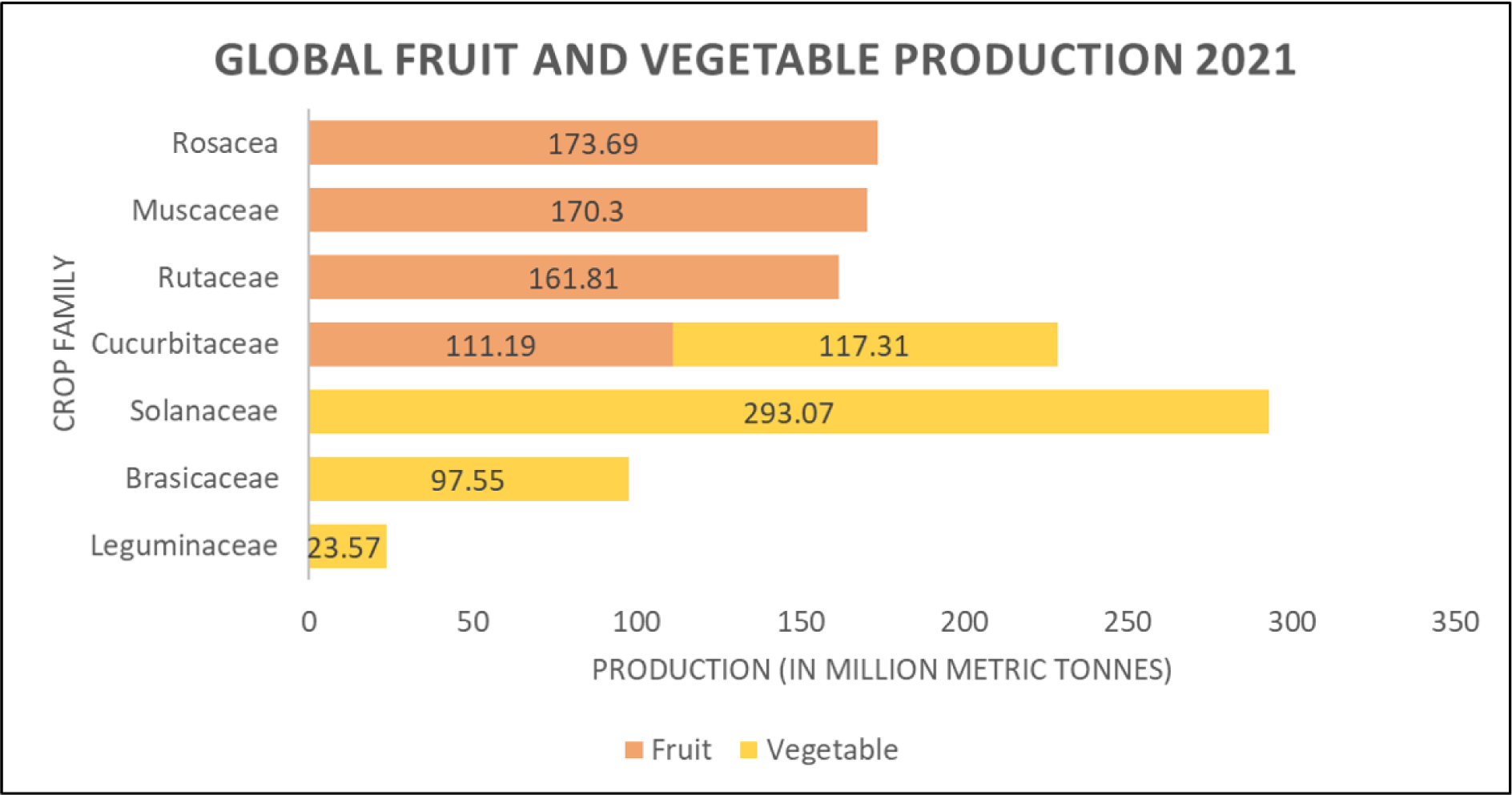
Global Fruit and Vegetable Production 2021 (Shahbandeh, M. (2023) (a & b)

**Fig 4.**
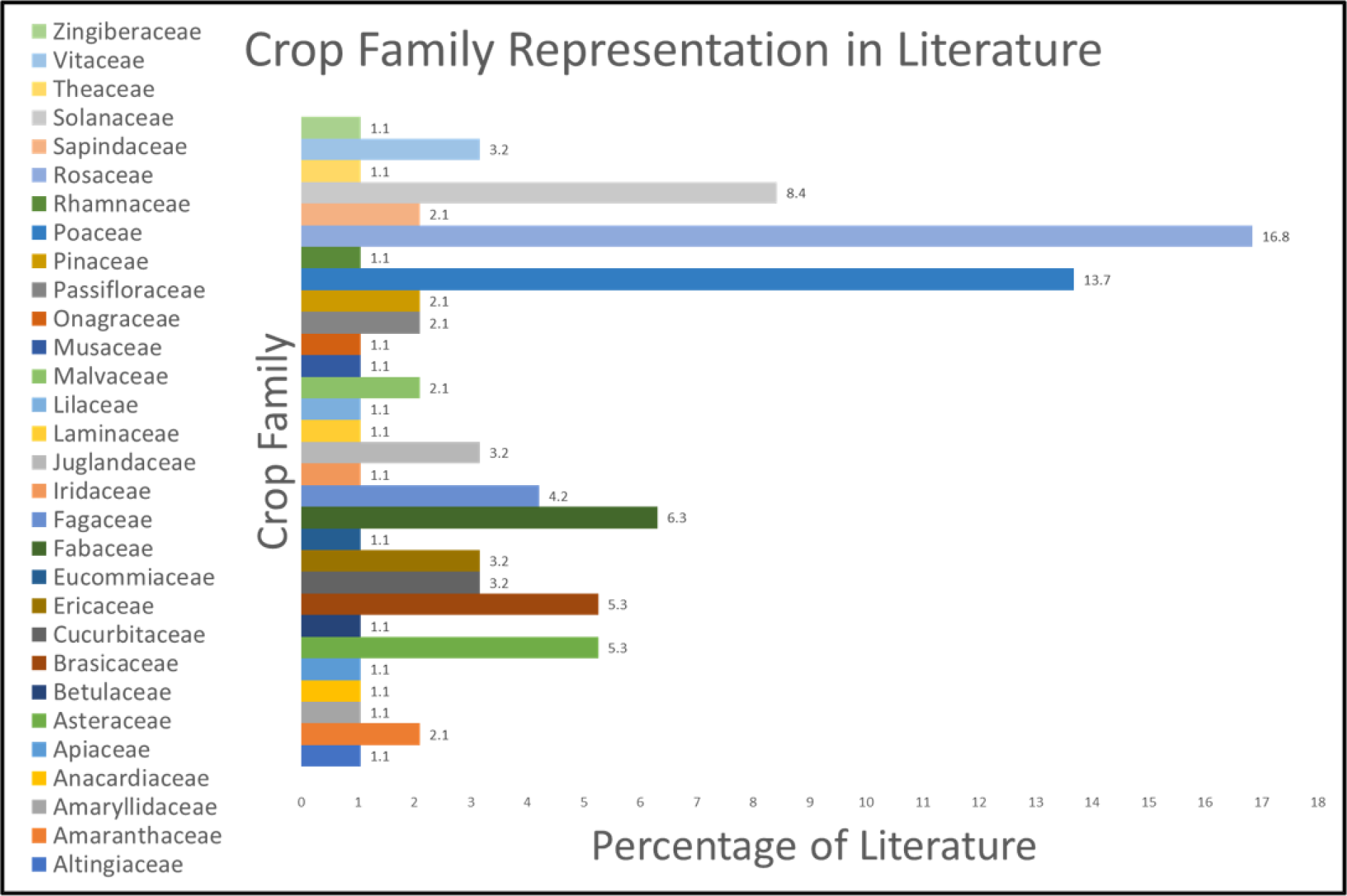
Crop family representation in UV reflectance literature from 1969 to 2021.

In agricultural remote sensing, the focal crop component indicates the physical parameter being researched. Fig. 5 illustrates the distribution of research across floral parts for our meta-analysis.

**Fig 5.**
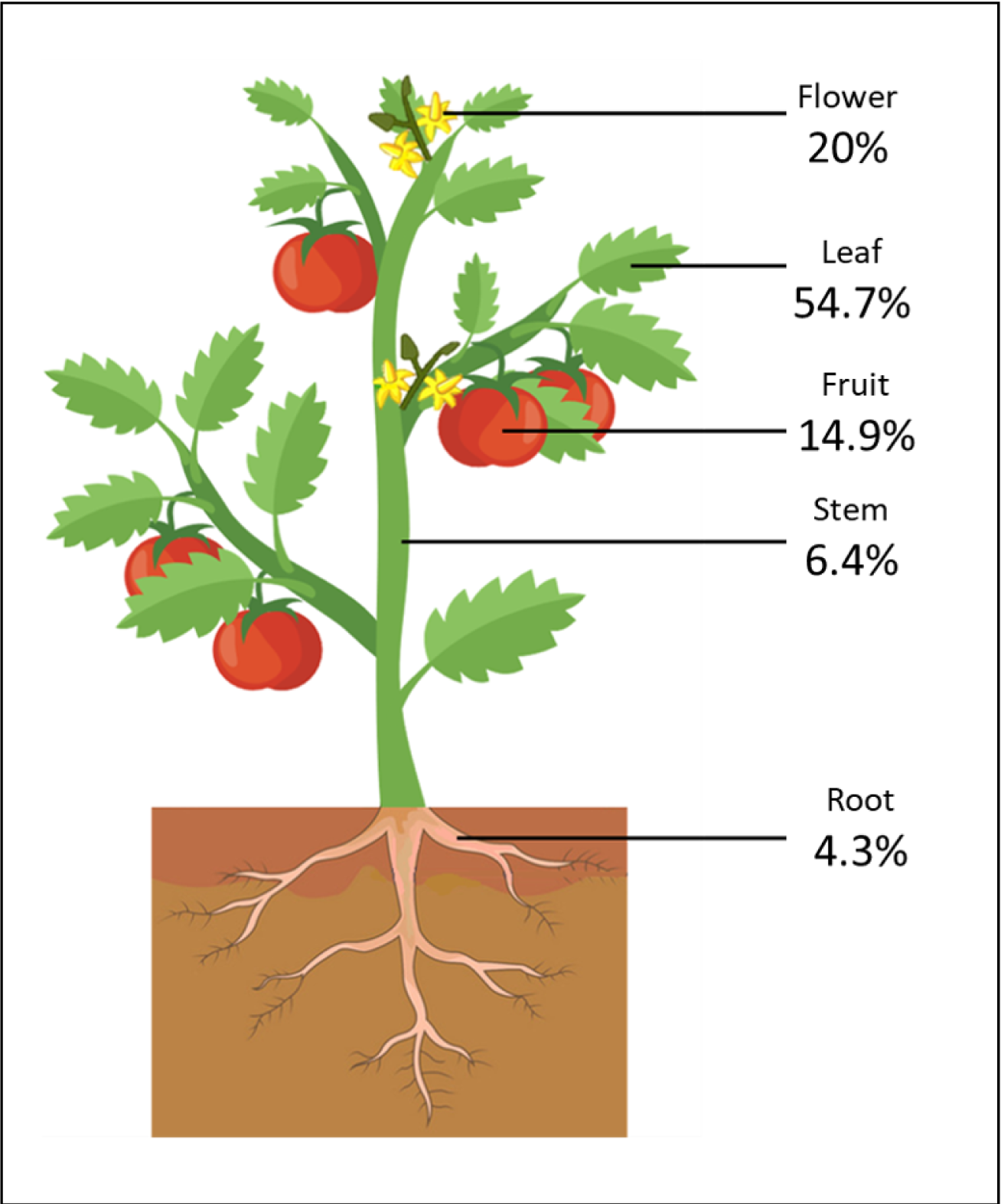
Floral parts spectrally analyzed in our literature meta-analysis.

We found that papers containing leaf reflectance represented most of the published research (55.7%), and these papers assessed growth rate, plant stress, or nutritional deficiencies. Publications focused on stems (6.4%) and roots (4.3%) assessed the quality of a given crop, e.g., lumber or tubers. Papers presenting the spectral reflectance of fruits (14.9%) had contents that varied the most, from assessing fruit ripeness and flavour quality to detecting disease or training detection algorithms for remote sensors. Papers analyzing flowers (20%) comprised two categories: pollinator-plant interaction and remote flower detection. However, it’s worth noting that over a third of all the flowers documented (31.57%, Tab. 1) were captured on UV film (Utech F.H. & Kawano S., 1975). Since UV film, like any camera film, is prone to human error during development, the reported reflectance pattern or intensity may not be accurate. For instance, Utech, F. H., & Kawano, S. (1975) reported a pattern of central petal absorption and UV reflecting anthers for two *Rosaceae* species *Fragaria x ananassa* ’Duchesne’ and *Rubus illecebrosus*’ Focke. However, spectrophotometric readings of wild *Fragaria* did not indicate this reflectance. Therefore, to fill the gap in the literature on strawberry (*Fragaia*) flower reflectance, one of the most significant contributors to the *Rosaceae* family’s global dominance in the fruit industry, we assessed the spectral reflectance of a variety of strawberry cultivars both quantitatively using a spectrophotometer and qualitatively using a UV sensitive camera.

### Spectral analysis of Strawberry cultivars

The *Rosacea* family contains many orchard species, such as apples and cherries, and berry species, such as strawberries and raspberries. Crops in the *Rosacea* family share similar floral phenotypic traits, such as five radially symmetrical sepals and petals, spirally arranged stamens, and a cup-like structure at the flower base known as a hypanthium (Folta, K. M.,& Gardiner,S.E.Eds.,2009). Due to their visual floral similarity, remote sensing and pollinator vision studies involving these crops tend to extend findings to the whole family (e.g., Dias P.A. et al., 2018; Eeraerts, M. et al., 2020). However, only some studies have investigated the actual spectral reflectance of *Rosacea* flowers.

### Methodology

#### Plants

Bare root plants of day-neutral *Fragaria ananassa sp.* cultivars (”Fort Laramie”, ”Hecker”, ”Seascape”), wild ancestor *Fragaria vesca*, and Asian *Fragaria ananassa x F. comarium* hybrid (”Berried Treasure Red”) were purchased from ©2020 Vesey Seeds. Plants were potted with a 2:2:1 ratio of acidic potting soil, shrimp compost, and sand in 7.5L containers and fertilized bi-monthly with 15-30-15 liquid feed. We removed flowers for imaging within 12 hours of opening and imaged the petal(P), anther(A), sepal(S) and upper leaf (L) from each flower. All plants used in this study were in good health and grown outdoors under natural light.

#### Reflectance spectra of Fragaria sp. flowers

We collected spectral reflectance measurements with a Perkin Elmer’s Lambda 850 UV-VIS spectrometer at the University of Laval in Quebec City, Canada. All flowers imaged were within 12 hours of first flowering and were intact. Each flower comprised three ’samples’: full flower upper side, petal only, and central anther and stamen disk only. We imaged the leaves of each cultivar on the upper and lower surfaces. At least two flowers or leaves per cultivar plant were measured. The Spectrophotometer was calibrated using Spectralon as suggested by the manufacturer. The measurement interval was set to 1nm with scans conducted over the 200-700nm range and repeated thrice per sample. Results were exported as an Excel spreadsheet of % reflectance values.

#### Quantifying contrast of floral parts

We quantified the visibility (*△*S)of strawberry flowers to pollinators using the Normalized Segment Classification (NSC) vision model (Rodŕıguez-Girońes, M. A., & Telles, F. J., 2020). Unlike previous segment classification models, the NSC model is 1) species independent and 2) considers brightness in its calculation. The model calculates a value (*△*S) based on the Euclidean distance between two spectrogram curves, indicating their contrast. The larger the number, the greater the contrast.

## Results

We present the spectrograms for *Fragaria vesca* (Fig. 6-a), the four *Fragaria x ananassa* cultivars (Fig. 6-b to d) and their respective leaves (Fig. 6-f) in Fig. 6(DOI 10.5281/zenodo.7847476). Table 2 presents the NSC vision model contrast values (*△*S) we obtained for each cultivar. We calculated *△*S values with leaves (L) and petals (P) to test floral contrast with leaf background. A floral pattern (e.g. bull’s eye pattern) was tested by comparing the outer floral part (petal, P) with the central floral part (anthers, s). We also tested sepals as they are visible when petals are damaged or a cultivar has sparse inflorescence.

**Fig 6.**
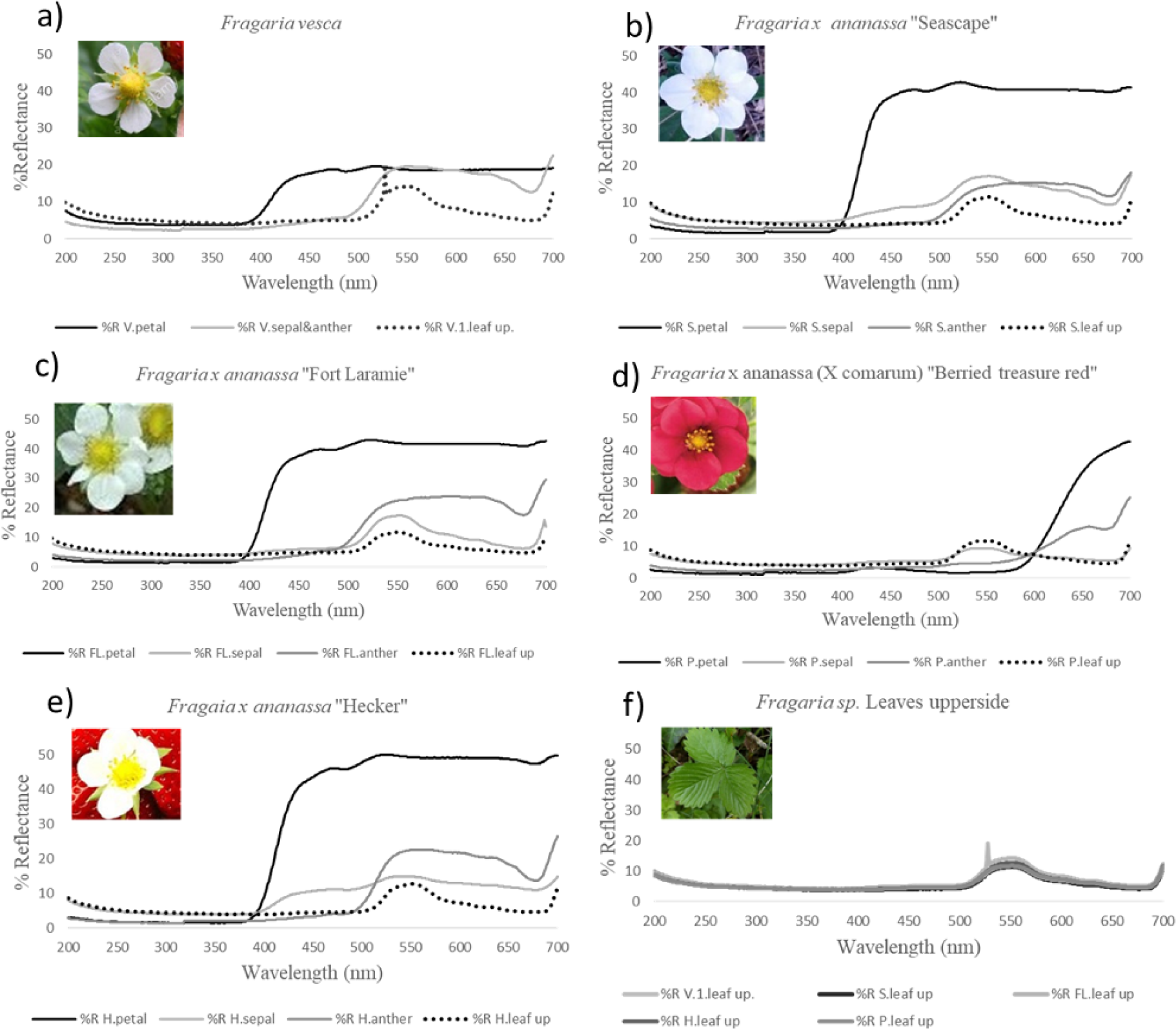
Reflectance spectrograms of Strawberry cultivars.

**Table 2.** Visibility of Fragaria sp. floral parts to trichromatic insect pollinators. *Indicates above bee contrast detection threshold.

| Cultivar | Floral parts contrasted | General trichromatic pollinator _S |
| --- | --- | --- |
| *Fragaria x ananassa (x comarum) 'Berried treasure Red'* | A/S | 1.165 |
|  | A/P | 0.255 |
|  | S/P | 1.309 |
|  | L/P | 1.335 |
| *Fragaria vesca* | A+S/P | *5.261 |
|  | L/P | *8.861 |
| *Fragaria x ananassa 'Hecker'* | A/S | 0.616 |
|  | A/P | *28.027 |
|  | S/P | *28.53 |
|  | L/P | *33.086 |
| *Fragaria x ananassa 'Fort Laramie'* | A/S | *3.663 |
|  | A/P | *21.001 |
|  | S/P | *24.651 |
|  | L/P | *27.127 |
| *Fragaria x ananassa 'Seascape'* | A/S | 2.204 |
|  | A/P | *24.638 |
|  | S/P | *22.441 |
|  | L/P | *27.463 |

### Leaves

Fig. 6-f demonstrates the minor variation in upper leaf reflectance across *Fragaria sp.* and cultivars indicating that the main factor in differing floral contrast and visibility to pollinators is solely the factor of floral pigments.

### Fragaria vesca

*Fragaria vesca* is a wild native strawberry specie that has well-documented pollinator-flower interaction (Blžytė-Čerěskienė, L. et al. 2012). Its spectrogram is published on the Floral Reflectance Database (FReD) and was used as a control for variation between studies. Our spectral reflectance curve showed the same pattern as previous reports (Chittka,et al., 1994). The flower shows a distinct flower petal peak in the Bee blue range (spectral peak 424nm, Skorupski, P., & Chittka, L. 2010) and a sepal/anther peak in the Bee green range (spectral peak 539nm, Skorupski, P., & Chittka, L. 2010). A study by Martinez-Harms, J. et al. (2010) found that bees could detect flowers 75% of the time with a contrast value (*△*S) as low as 2.3. The contrast value for petal (*△*S=8.861) and sepal/anther (*△*S= 5.261) support the observations made in previous behavioural studies that this flower is visible to pollinators.

### Fragaria x ananassa (x comarium) ’Berried treasure Red’

The red flowering cultivar, *Fragaria x ananassa (x comarium)* ’Berried treasure Red’ (Fig. 6-d), had floral peaks beyond 600nm and showed no contrast values between floral parts *△*S *>* 1.34, indicating that bee pollinators would be blind to this cultivar. This cultivar, in particular, was bred purely for aesthetic appeal, with little regard for yield potential. It is not cultivated in fields and is a newer release to the consumer market.

### White-flowering cultivars

When compared to their wild counterpart (Fig. 6-a), the white flowering cultivars ”Seascape,” ”Fort Laramie,” and ”Hecker” (Fig 6, b, c, and e, respectively) exhibit higher petal reflectance, creating higher contrast and visibility for pollinators. These cultivars demonstrate petal peaks in the bee blue and anther peaks in bee Green (Spectral peak 539 nm, Skorupski, P., & Chittka, L. 2010). Anther/ petal contrast values indicate a discernable Bull’s eye pattern for all three cultivars in the bee blue/bee green range (*△*S= 24.638, 24.651, and 28.027, respectively). The highest contrast value for all three cultivars was between petals and background leaves (*△*S= 27.463, 27.127, and 33.086, respectively). ”Hecker” had the highest contrast value with an L/P *∼*S = 33.086; 26.8% higher than its wild counterpart. ”Hecker” is noted for producing large berries with good flavour (Bringhurst, R., & Voth, V., 1980). We know that insect pollination has a direct, positive effect on fruit quality (e.g., Nye, W. P., & Anderson, J. L.,1974; Wietzke, A., et al. 2018; Abrol, D. P., et al. 2019). In selectively breeding for higher yield and better-quality fruit, breeders could have inadvertently selected more visible flowers. As such, ”Hecker” flowers would be more visible from the air to nearby pollinators than their conspecifics. Bees preferentially visit more visible targets when nectar rewards are equal (Spaethe, J. et al. 2001). The higher visitation rate by natural pollinators would positively affect the outcome of yield assessments.

### Strawberry flowers in the UV

Gyan, K. Y. & Woodell, S. R. J. (1987) documented the spectrogram of the blackberry, *Rubus fructicosus*, which indicated a *∼* 35% reflectance around 360nm. Almond cultivars, *Prunus dulcis*, have also shown a consistent, distinct peak at 350nm (Chen, B., Jin, Y., & Brown, P., 2019). Although the *Fragaria* genus shares the same family as Rubus, Fig. 7 does not indicate any reflectance peaks around 350-360 nm (or Bee UV, spectral peak 347nm, Skorupski, P., & Chittka, L. 2010) across all study species. Incidentally, there is minimal spectral reflectance in the near UV range (300-400nm). However, all the above samples exhibit an increase in reflection below 250nm. In this way, the *Rosacea* family shows reflection diversity in the near UV spectrum, indicating that findings from remote sensing or pollinator vision studies should not extend beyond the specie or cultivar at hand.

## Conclusion

At this time, future floral reflectance studies should put more emphasis on crop species than native species. Our results showed a need for lightweight camera models for in-situ UV remote sensing. Current models are costly and cumbersome for automated deployment. The representation of crop families in the literature reflects their economic value in the global market. That being said, UV reflectance is still a tiny proportion of all crop spectral reflectance studies. To add to the public database of crop spectral reflectance, we spectrophotometrically analyzed five strawberry cultivars (DOI 10.5281/zenodo.7847476). We studied the data and noted that commercial white-flowering strawberries produced a bull’s eye contrast pattern in the bee-blue/ bee-green, producing the highest contrast with background leaves. The most notable, ”Hecker,” is prized for its production volume and high fruit quality. This may be due to its greater visibility to pollinators, i.e. bees, leading to higher pollination rates. Further studies presenting the spectral reflectance of crops across pollinator vision range (near UV to blue) would benefit pollinator interaction research and the agricultural industry and be an excellent resource for crop breeders.

## Acknowledgements

We would like to thanks NSERC for funding this study. Our deepest thanks to Marie-Hélène Forget and the biology department at Laval University, Qc. for hosting us on their campus and for the loan of their lab equipment. Without both, this study would not have been possible.

